# CovidOutcome2: a tool for SARS-CoV2 mutation identification and for disease severity prediction

**DOI:** 10.1101/2022.07.01.496571

**Authors:** Regina Kalcsevszki, András Horváth, Balázs Győrffy, Sándor Pongor, Balázs Ligeti

## Abstract

Our goal was to develop a platform, CovidOutcome2, capable of predicting disease severity from viral mutation profiles using automated machine learning (autoML) and deep neural networks applied to the available large corpus of sequenced SARS-CoV2 genomes. CovidOutcome2 accepts either user-submitted genomes or user defined mutation combinations as the input. The output is a predicted severity score plus a list of identified, annotated mutations and their functional effects in VCF format. The best model performance is a ROC-AUC 0.899 for the model including patient age and ROC-AUC 0.83 for the model without patient age.

**Availability:** CovidOutcome is freely available online under the URL https://www.covidoutcome.bio-ml.com as well as in a standalone version https://github.com/bio-apps/covid-outcome.

## Introduction

The number of deposited SARS-COV-2 genome data is unprecedentedly large: as of April 2022 more than 10 million COVID genomes were deposited in the GISAID database (1) and it opened the way for the application of machine learning algorithms. One of the most important questions is how virus mutations are related to the severity of the disease and if disease severity can be predicted from genomic mutations. Previously we applied autoML to COVID amino acid replacements (2). Here we present a new tool, CovidOutcome2 that uses deep learning neural networks in addition to autoML on an approximately 8-fold larger training set. Moreover, it offers a new pipeline for evaluating user defined viral variant combinations, multiple alignment based mutation calling and functional predictions.

## Data and methods

First, we retrieved the 67710 SARS-CoV-2 genomes with the corresponding patient and annotation data from the GISAID database (https://www.gisaid.org/, accessed on March 18, 2022). Two cohorts were defined “mild” and “severe” based on the normalized patient status field (supplementary table 1). Two balanced, stratified (by age groups and time periods) training datasets were compiled, one with additional age information (dataset 1, prefix: CA) and one without it (dataset 2, prefix C; supplementary table 2, and supplementary table 3). The age attribute was standardized and cleaned (more detailed description in supplementary file 1). The NCBI SARS-CoV-2 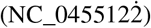 genome was used as a reference genome in the quality control and in the mutation detection parts of the pipeline. Sequences that do not match the prerequisites were discarded. For more details about the QC process see the supplement. The mutation calling (SNVs and indels) is based on the multiple alignment (calculated by MAFFT (3) containing the reference genome. Large insertions and deletions (length > 50) were filtered out, as well as the low frequency (present <10% either in mild or severe group) and synonym mutations.

We applied two different machine learning strategies: i) autoML and ii) a deep, fully-connected neural network. The autoML tool JadBio (4) performs the following steps, i) it analyzes the uploaded data and identifies the list of algorithms and applicable feature selection methods, along with their hyperparameters; ii) it generates the possible configurations (preprocessing + feature selection + predictive model) iii) it evaluates the configurations using 20 repeated 10 fold CV and identifies the best model, iv) it estimates the model performance using BBC-CV (5). In our case a regularized logistic regression model combined with feature selection based on Statistically Equivalent Signatures (SES) was found optimal. We also developed a fully connected, multilayer neural network consisting of 7 linear layers with ReLU activation functions. The number of neurons in each layer was {251,256,256,256,128,64,32}. The models were trained with batch size 128, via 500 epoch, binary cross entropy with logit loss were optimized with Adam optimizer and were implemented using PyTorch library (https://pytorch.org).

## Results and discussions

CovidOutcome offers two pipelines for predicting disease severity starting either from i) a user-defined mutation combination, or ii) from a user-submitted genome (Figure 1). In the first pipeline the user can upload up to 100 COVID related sequence data in either fasta format or plain text. The webserver first performs a sequence quality assessment (i.e. number of N characters, substantial similarity to the reference genome, etc.), the genomic mutations are then called, annotated by SNPEffv5 (6). The identified mutations are filtered (i.e. synonym variants are removed) and finally the mutation profile is created and the prediction is carried out. The user can optionally specify the age of the patient for each sample. In cases where no age information is available a model trained without age information is used for the prediction. The results page presents the identified mutations with their annotations, their predicted effects and their frequencies, as well as the prediction score and categories for the outcome (i.e. confident-mild, confident-severe, etc). The outputs, i.e. mutation annotations, prediction scores and categories can be downloaded in standard formats, such as VCF, for further evaluation.

**Fig. 1.**
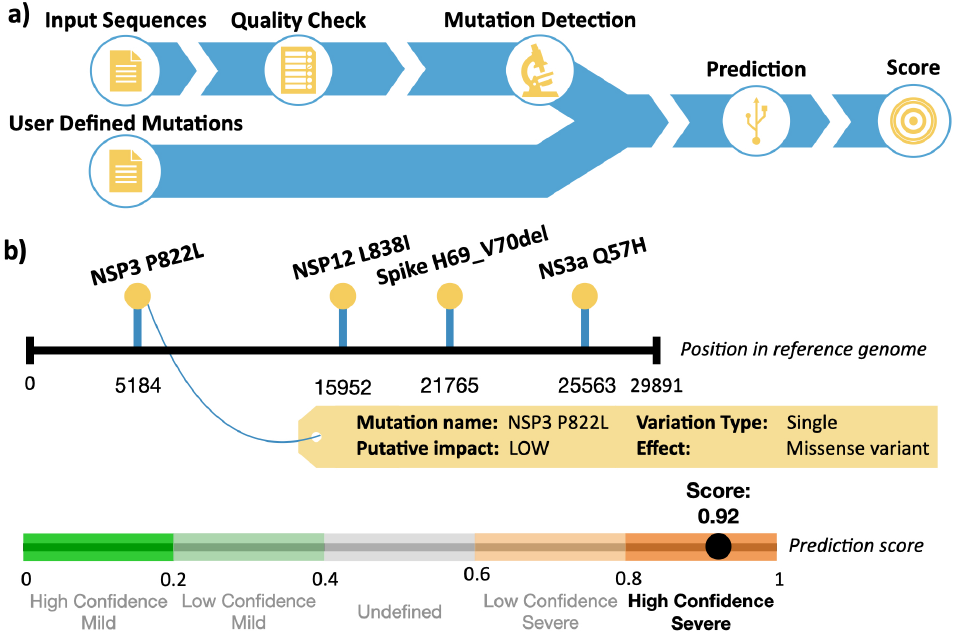
CovidOutcome pipelines. a) Pipelines for prediction from genomic sequences (i) and user/defined mutation lists. b) Outputs with graphical presentation of mutations and a severity score with prediction confidence (more in supplementary file 1)

The inputs of the second pipeline are the user defined covid-variants which can be not only nucleotide, but also protein mutations. It is possible to add age information for the newly defined virtual genomes and the user can choose from two different types of ML approaches similarly to the previous pipeline. The prediction scores and the detailed information about the mutations are presented on the results page.

Model performances were evaluated using 10 repeated 10 fold cross validation, which is summarized in **Table 1**. The best performing models were provided by the neural network (with ROC-AUC=0.89 (dataset 1) and 0.83 (dataset 2). JADBio identified the Statistically Equivalent Signature (SES) algorithm (with hyper-parameters: *maxK* = 2, *α* = 0.05 and budget = 3 * *nvars*) the best feature selection method and Ridge Logistic Regression (with penalty hyper-parameter *λ* = 1.0) as classification approach. For a more detailed description see the wiki page of CovidOutcome as well as for the list of identified mutation signatures. (https://github.com/bio-apps/covid-outcome/wiki/CovidOutcome).

**Table 1.**
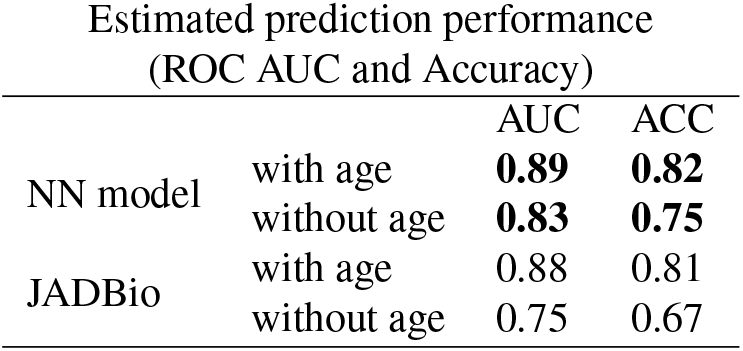
The estimated model performances using 10-repeated 10 fold cross validation. The applied metrics are ROC AUC (Area under the ROC curve and prediction accuracy.)

| Estimated prediction performance<br>(ROC AUC and Accuracy) |  |  |  |
| --- | --- | --- | --- |
|  |  | AUC | ACC |
| NN model | with age | **0.89** | **0.82** |
|  | without age | **0.83** | **0.75** |
| JADBio | with age | 0.88 | 0.81 |
|  | without age | 0.75 | 0.67 |

As any other machine learning technique, our models are also limited to the data they were trained on. The current model is unable to predict the potential effect of completely new mutations that do not exist at least once in GISAID. It is well-known that the COVID severity depends on many factors (i.e. disease comorbidity, healthcare-service, etc.), which are naturally not included in our model, except the age of the patient. We believe that frequently updating the models with new samples could at least partly resolve the above mentioned problems.

## Conclusion

In this work we presented an extended tool and service for investigating the association between the SARS-CoV-2 viral mutation signatures and disease outcome using a combination of machine learning and bioinformatics techniques. CovidOutcome2 leverages the large amount of public genomic data and offers two pipelines for predicting the possible outcome of the disease. The updated data representation integrates not only the substitutions, but indels and relevant UTR mutations as well. The output is a disease severity score as well as a list of mutations in a standardized format VCF.

## ACKNOWLEDGEMENTS

The authors wish to acknowledge the support of ELIXIR Hungary (www.elixir-hungary.org) and Dr. I. Tsamardinos (University of Crete, Greece) for help and advice.

