## Supplementary file 1 for "CovidOutcome2: a tool for SARS-CoV2 mutation identification and for disease severity prediction"

June 2022

#### 1 Sequence data, preprocessing and mutation calling

##### 1.1 Training sets

First, we retrieved the 67710 SARS-CoV-2 genomes with the corresponding patient and annotation data from the GISAID database (https://www.gisaid.org/, accessed on March 18, 2022). The patient status field contains submitter defined terms and categories therefore standardization was needed. We defined two cohorts: "mild" and "severe". The age was standardized and cleaned. The records with undefined value were discarded. The number of mild samples are considerably larger than the severe samples. To avoid the genomic mutations overrepresented in mild samples we created a balanced dataset, which contains equal number of mild and severe samples. We also stratified the sets to time periods (quarters of the years), beacause the severe-mild ratio greatly varied. However, the viral properties are driven by it's genomic content, large number of studies confirmed that the disease outcome and susceptibility of COVID-19 is statically linked to various comorbidities, age, and directly or indirectly to the mutations in the virus genome. To support analyses where the age is unkown, we created a training dataset stratified to age groups. We compiled two balanced, stratified training datasets, one with additional age information (dataset 1, prefix: CA) and one without it (dataset 2, prefix C; supplementary file 1, and supplementary file 2).

##### 1.2 Sequence data preproccessing and mutation calling

The NCBI SARS-CoV-2 (NC_0455122̇) genome was used as a reference genome in the quality control and in the mutation detection parts of the pipeline. The sequence IDs are replaced with unique IDs. Sequences that do not match the following prerequisites were filtered out: a) The length of the sequences: 26000-33000 base pairs; b) the ACGT ratio of the sequences is higher than 0.95; c) the congruence score of the alignment to the reference genome is higher than 0.75. The congruence score is calculated based on a Mummer alignment of the sequences to the reference genome. The congruence score is calculated as 1 - (D + F) / L, where D is the number of different nucleotides from the reference, F is how many more bases the sequence has than the reference, and L is the length of the reference. The mutation calling starts with running MAFFT [Katoh et al., 2002] to align the sequences in the fasta files with the reference sequence. Based on the alignment, SNVs and indels are called for each sample, then subsequent mutations are grouped together. Large insertions and deletions (length $> 50$) were filtered out, as well as the low frequency (present $< 10\%$ either in mild or severe group) and synonym mutations.

### 2 Pipelines

#### 2.1 Pipeline for outcome prediction from genomic data

In this pipeline the user can upload up to 100 COVID related sequence data in either fasta format or plain text. The webserver processes and analizes the sequences figure 1). It performs a quality assessment (i.e. number of N characters, substantial similiarity to the reference genome), followed by multiple sequence alignment steps. ( The genomic mutations are called based on the MSA and mutation effects are predicted and annotated by SNPEffv5 [Cingolani et al., 2012]. Then the identified mutations are filtered (i.e. synonym variants are removed) and finally the mutation profile is created and the prediction is carried out. The user can optionally specify the age of the patient for each sample. In cases where no age information is available a model trained without age information is used for the prediction. The pipeline offers two types of ML approaches for the prediction a) using deep neural network model, b) applying a traditional, interpretable model (feature selection – Test-Budgeted Statistically Equivalent Signature (SES) algorithm – with regularized logistic regression model) optimized by JadBIO autoML platform [Tsamardinos et al., 2020]. The results page presents the identified mutations with their annotations, their predicted effects and their frequencies, as well as the prediction score and categories for the outcome (i.e. confident-mild, confident-severe, etc). The outputs, i.e. mutation annotations, prediction scores and categories can be downloaded in standard formats, such as VCF, for further evalutation.

#### 2.2 Pipeline for outcome prediction of user defined viral variants

Our recently added pipeline's goal was to give prediction about the expected outcome for novel viral variants, even for non-existing ones based on the accumulated mutation data deposited in GISAID. The inputs of the pipeline are the

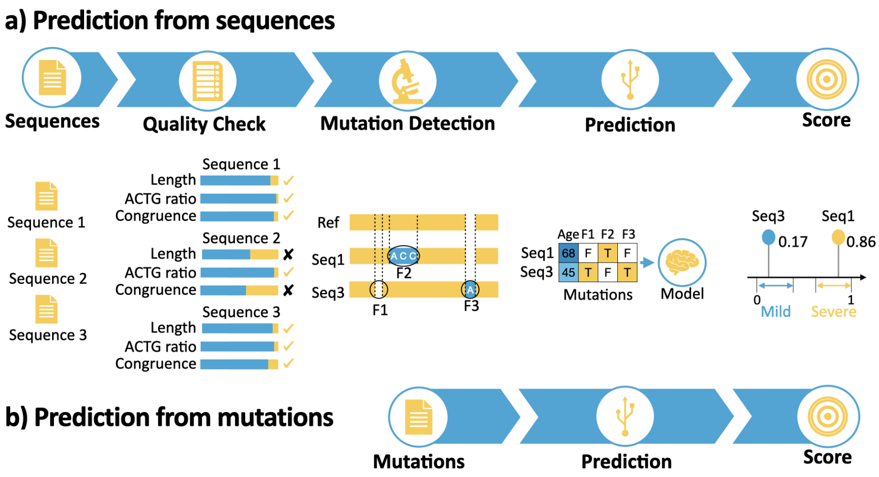


Figure 1: **CovidOutcome pipelines**. a) Pipeline for prediction from variants. b) Pipeline for outcome prediction from genomic data.

user defined covid-variants which can be not only nucleotid, but protein mutations (indels, insertion) as well, coded as standard. One of the main limitations of the pipeline is that it is only 'aware' of the currently known mutations, therefore it is unreliable with unseen or completely new mutations. In this pipeline the user can define multiple virtual genomes or SARS-CoV2 variants. It is possible to add age information for the newly defined virtual genomes and the user can choose from two different types of ML approaches similarly to the previous pipeline. The prediction scores and the detailed information about the mutations are presented on the results page.

### 3 Outputs

After uploading the input sequences to the webserver and choosing the prediction model, the prediction pipeline is run and three output table are presented in the result page: the Sequence table, the Mutation table and the ML table. Each table can be downloaded in *tsv* format.

#### 3.1 Sequence table.

The input sequences go through a thorough quality check before mutation detection. *Sequence table* contains information about the quality measures of each input sequence. The columns of the table are the following:

**User ID:** Sequence ID given by the user input.

**Sequence Length:** Length of the sequence.

**Sequence Header:** Fasta header of the sequence.
**Sequence ID:** Unique sequence ID given by CovidOutcome pipeline.
**Ids:** OK, if Sequence ID is given.
**Uppercase:** OK, if the sequence is changed to uppercase characters.
**Sequence Length:** OK, if the sequence length is appropriate (default: 0-35000).
**ACTG Proportion:** OK, if the ratio of ACTG characters in the sequence is high enough (default: 0.95).
**GC Content:** OK, if the GC content of the sequence is appropriate (default: 0-100).
**Congruence:** OK, if the congruence score of the sequence is high enough (default: 0.75).
**Summary:** OK, if all checks are OK.

#### 3.2 Mutation table.

*Mutation table* contains the annotated mutations that were found in the input sequences. The columns (#CDS Position / Length, Distance to feature, Up/Downstream, Intergenic, Effect, Gene Name, HGVS.p, ID, Protein Position / Length, Putative impact, Transcript biotype, Variation Type, cDNA Position / Length, Note) are provided by SNPEffv5 [Cingolani et al., 2012]. The descriptions of the additional columns are the following:
**Mutation Length:** Length of the mutation.
**Mutation name:** Mutation name in standard form. In case of coding regions: {protein name}_{mutation in HGVS format}. In case of UTR regions: {protein name before}-{protein name after}_n.{mutation in HGVS format}.
**Protein Name:** Name of the affected protein.
**Sequence ID:** Unique sequence ID given by CovidOutcome pipeline.
**User Sequence ID:** Sequence ID given by the user input

#### 3.3 ML table.

The model's features are selected from the annotated mutations and the model is evaluated on the samples, resulting in a prediction for the disease outcome. *ML table* contains information about the prediction for each sample. The columns of the table are the following:
**User ID:** Sequence ID given by the user input.
**Age:** Age information if it is given by the user.
**Mild Prediction Score:** Probability of whether the outcome is mild.
**Prediction Score:** Probability of whether the outcome is severe.
**Applied Model Type:** Type of model used for prediction: *deep_model_with_age*. *deep_model_without_age*, *automl_model_with_age*, *automl_model_without_age*
**Prediction Confidence:** Confidence of the prediction: *high confidence* (0-0.2 and 0.8-1), *low confidence* (0.2-0.4 and 0.6-0.8), *undefined* (0.4-0.6).
**Label:** Predicted label for the sample: *mild* (0-0.5), *severe* (0.5-1).

Kazutaka Katoh, Kazuharu Misawa, Kei-ichi Kuma, and Takashi Miyata. Mafft: a novel method for rapid multiple sequence alignment based on fast fourier transform, 2002.

Ioannis Tsamardinos, Paulos Charonyktakis, Kleanthi Lakiotaki, Giorgos Borboudakis, Jean Claude Zenklusen, Hartmut Juhl, Ekaterini Chatzaki, and Vincenzo Lagani. Just add data: Automated predictive modeling and biosignature discovery, 2020.
